# B vitamin quantification in lentil seed tissues using ultra-performance liquid chromatography-selected reaction monitoring mass spectrometry

**DOI:** 10.1101/2022.12.07.519471

**Authors:** Jeremy Marshall, Ana Vargas, Kirstin Bett

## Abstract

Lentils are an important source of macronutrients, including protein and fiber, as well as micronutrients such as vitamins and minerals, especially in a plant-based diet. Quantifying variation among genotypes, including wild germplasm, is desirable to better understand the genetics of differential B vitamins content for breeding of this trait and to understand their potential contributions to the lentil crop. We analyzed thirty-four cultivated and three wild genotypes for vitamins B1, B2, B3, B5, B6, B7, and B9. Seeds were assayed whole, and separated into cotyledons only, or seed coats only. Variation for all B vitamins was observed across the cultivars. Overall, cotyledons had higher concentrations of B1 and B3, while seed coats had higher concentrations of B2, B5, B6, and B9. Wild accessions had the highest concentrations of vitamin B9 and were also among the highest for vitamin B2. These results demonstrate the differential distribution of B vitamins across seed tissues and lentil genotypes, and that dehulling prior to consumption results in the loss of B vitamins otherwise available in whole seeds. They also indicate there is genetic variability which could be used to increase B-vitamin levels in lentil via breeding.

## 1. Introduction

Pulse crops continue to garner more interest globally as sources of protein, fiber, and various other nutrients. The high protein content of many pulses has made them particularly attractive to the plant-based protein industry, which has expanded in recent years (NRC, 2019). From 2010 to 2017 sales of plant-based meat alternatives increased by an average of 8% annually, with Canadian sales during fiscal year 2016/2017 exceeding $1.5 billion (NRC, 2019). Lentils are also high in other nutrients, are flexible in cooking, and easy to prepare (Khazaei, *et al*., 2019; Kumar *et al*., 2016).

Lentils are known for their nitrogen fixing ability, making them popular choices during crop rotation to help maintain soil quality (Lupwaky and Kennedy 2007). In 2019, global production of lentils was 5.73 million tonnes (FAO, 2019). Canada is the global leader in lentil production (2.17 million tonnes 2019), followed by India (1.23 million tonnes 2019) and Australia (0.53 million tonnes 2019) (FAO, 2019).

With this increased interest in pulse consumption, obtaining accurate information on individual micronutrient concentrations across pulse species can aid in the selection of optimal crops for select nutritional benefits. A recent review (Hall *et al*., 2017) shows the variation in micronutrient concentrations among genotypes of the same species across pulse crops, suggesting that characterization of nutrient profiles may contribute to the identification of genotypes of interest for breeding.

The review by Hall *et al*. also demonstrates gaps in data collection for certain micronutrients, including B vitamins. B vitamins are a family of eight micronutrients which are essential vitamins in the human diet and function as cofactors for numerous metabolic pathways. The B vitamin families are: B1 thiamine, B2 riboflavin, B3 nicotinic acid, B5 pantothenic acid, B6 pyridoxal, B7 biotin, B9 folate, and B12 cobalamins. All the B vitamins, other than vitamin B12, are produced in plants, and can be obtained through plant-based dietary sources. Deficiencies in B vitamins can lead to a number of health-related illnesses including cardiovascular or neurological conditions of beriberi (B1), cancer and cardiovascular or neurological conditions (B2), dermatitis and neurological conditions (B3 and B6), as well as cardiovascular disease and impaired fetal neural tube development (B9) (Baj and Sieniawska, 2017).

Only the various forms of B9 have been extensively examined in pulses, with limited examination of B1 and B2 present in a small number of pulse species (Hall *et al*., 2017). This leaves many B vitamins largely uncharacterized, which is a gap in knowledge that can and should be resolved.

Previously, a B vitamin extraction and liquid-chromatography coupled to mass-spectrometry analysis was optimized, that allows for the quantification of sixteen distinct vitamers from the seven B vitamin families present in plants; one for B1, one for B2, two for B3, one for B5, three for B6, one for B7, and seven for B9 (Zhang *et al*., 2021). This analysis method allows for high-throughput analysis of pulse crop samples and quantification of all B vitamin families.

Our objective in this study was to use the previously optimized B vitamin analysis protocol to determine the complete B vitamin profiles for thirty-seven lentil genotypes - thirty-four cultivars and three wild accessions. As wild lentils possess genes of interest for breeding cultivated lines, including those associated with disease resistance or stress tolerance (Gela *et al*., 2021; Gorim and Vandenberg 2017), we sought to determine if there were any vitamin profile differences between the cultivated and wild accessions, we should be aware of. In addition, the analyzed samples were separated into whole seed, cotyledons only, and seed coats only samples, allowing the quantification of vitamin distribution across seed tissues. As split lentils, which have had their seed coats removed, are a commonly used pulse in recipes, we wanted to determine the impact on vitamins present following the loss of seed coat. The results of these analyses can be used to inform breeding line selection when seeking to maximize B vitamin concentrations without fortification.

## 2. Materials and Methods

### 2.1 Plant material

Lentil genotypes analyzed in this study listed in **Table 1** were grown in Saskatchewan, Canada. Three cultivars (CDC Greenstar, Eston, and Lupa) and three wild accessions (IG 72643, IG 72623, BGE 016880) were grown in Sutherland (52°09’58.2” N, 106°30’21.8” W) and Preston (52°07’23.9” N, 106°37’11.9” W) in 2018 and Sutherland in 2020, and all thirty-four cultivars were grown in Sutherland in 2020. The plants were grown in single plots consisting of three rows (1 m long) with an interrow spacing of 30 cm and a target population of 60 plants per plot. The soils are loam at all locations. Plots were grown under rainfed conditions with a total in-season rainfall of 140.3 mm in 2018 and 240.9 mm in 2020 (Environment Canada https://climate.weather.gc.ca/historical_data/search_historic_data_e.html).

**Table 1.** Lentil accessions analyzed for their B vitamins concentration.

| Name | Species | Cotyledon Colour | Origin | Source |
| --- | --- | --- | --- | --- |
| <u>Cultivars</u> |  |  |  |  |
| CDC Redberry | *Lens culinaris* | red | Canada | CDC-U of S<sup>β</sup> |
| CDC Robin | *Lens culinaris* | red | Canada | CDC-U of S |
| ILL 11551 | *Lens culinaris* | red | India | ICARDA<sup>ε</sup> |
| ILL 1704-1 | *Lens culinaris* | red | Ethiopia | ICARDA |
| ILL 213 | *Lens culinaris* | red | Afghanistan | ICARDA |
| ILL 4609 | *Lens culinaris* | red | Pakistan | ICARDA |
| ILL 5722 | *Lens culinaris* | red | Turkey | ICARDA |
| ILL 618 | *Lens culinaris* | red | Tajikistan | ICARDA |
| ILL 7727 | *Lens culinaris* | red | Morocco | ICARDA |
| LR-59-81<sup>∞</sup> | *Lens culinaris* | red | breeding line |  |
| PI 182217 | *Lens culinaris* | red | Turkey | USDA-ARS<sup>α</sup> |
| Ranjan | *Lens culinaris* | red | India |  |
| 02M-12 | *Lens culinaris* | red | breeding line | CDC-U of S |
| 2275-15 | *Lens culinaris* | red | Canada |  |
| 3954-105 | *Lens culinaris* | red | Canada | CDC-U of S |
| 4656-3 | *Lens culinaris* | red | Canada | CDC-U of S |
| CDC QG-1 | *Lens culinaris* | green | Canada | CDC-U of S |
| CDC Greenstar | *Lens culinaris* | yellow | Canada | CDC-U of S |
| CDC Milestone | *Lens culinaris* | yellow | Canada | CDC-U of S |
| Eston | *Lens culinaris* | yellow | Canada | CDC-U of S |
| Indianhead | *Lens culinaris* | yellow | Canada | CDC-U of S |
| Lupa | *Lens culinaris* | yellow | Spain |  |
| VIR 421 | *Lens culinaris* | yellow | Russia |  |
| Guarena | *Lens culinaris* | yellow | Spain |  |
| IG 858 | *Lens culinaris* | yellow | Algeria | ICARDA |
| ILL 1959 | *Lens culinaris* | yellow | Ethiopia | ICARDA |
| ILL 8007 | *Lens culinaris* | yellow | breeding line | Bangladesh |
| ILL 9 | *Lens culinaris* | yellow | Jordan | ICARDA |
| ILL 927 | *Lens culinaris* | yellow | Turkey | ICARDA |
| PI 431884 | *Lens culinaris* | yellow | Iran | USDA-ARS |
| PI 432005 | *Lens culinaris* | yellow | Iran | USDA-ARS |
| PI 432245 | *Lens culinaris* | yellow | Lebanon | USDA-ARS |
| Shasta | *Lens culinaris* | yellow | USA | USDA-ARS |
| 5365-6 | *Lens culinaris* | yellow | breeding line |  |
| <u>Wild Accessions</u> |  |  |  |  |
| BGE 016880 | *Lens orientalis* | red | Israel |  |
| IG 72643 | *Lens orientalis* | red | Syria |  |
| IG 72623 | *Lens odemensis* | red | Turkey |  |

<sup>∞</sup>LR-59-81: interspecific line (Eston *L. culinaris* × L01-827A *L. ervoides*), CDC, University of Saskatchewan, Saskatchewan, Canada. <sup>β</sup>CDC- U of S: Crop Development Centre, University of Saskatchewan. <sup>ε</sup>ICARDA: International Center for Agricultural Research in the Dry Areas.
<sup>α</sup>USDA-ARS: US Department of Agriculture-Agricultural Research Service.

Whole seeds were collected for each genotype at maturity and air-dried at 45°C until a moisture <13% was reached. Subsets of dried seeds were separated into cotyledons and seed coats using a Satake huller TM05C(2) (Stafford, TX, USA). Whole seeds, cotyledons only, and seed coats only, were ground to a fine powder using an Udy grinder (Fort Collins, CO, USA). All samples were aliquoted and stored at –80 °C until analysis. Analysis of the 2020 lentil samples included thirty-seven genotypes, with thirty-four cultivars and three wild accessions (**Table 1**). Samples included whole seeds, cotyledons only, and seed coats only for thirty-two cultivars, while two cultivars (Guarena and ILL 11551) and the wild accessions (IG 72643, IG72623, BGE 016880) had cotyledons only and seed coats only due to limited seed amounts. Genotypes harvested in 2018 were analyzed 25 months and the set grown in 2020 was analyzed 13-15 months, after harvest.

### 2.2 Chemical reagents and enzymes

2-Mercaptoethanol (MCE), 2-(N-morpholino)ethanesulfonic acid (MES), sodium ascorbate, acid phosphatase (AP), β-glucosidase from almonds (BGL) and activated charcoal were purchased from Sigma-Aldrich (St. Louis, MO, USA). Dithiothreitol (DTT), potassium phosphate monobasic and dibasic, ammonium acetate and acetic acid were purchased from Fisher Scientific (Fair Lawn, NJ, USA). LC-MS grade acetonitrile and LC-MS grade methanol (MeOH) were purchased from Thermo Fisher Scientific (Nepean, ON, Canada). Fresh rat serum (Lampire Biological Laboratories, Pipersville, PA, USA) was shipped on wet ice, immediately aliquoted into 10 mL stocks upon arrival, and stored at –80 °C until needed. The AP and BGL were prepared in water to final concentrations of 5 mg/mL. Endogenous B vitamins in the rat serum were removed by adding one-tenth (w/v) of activated charcoal (Jha *et al*., 2015; Zhang *et al*., 2018a, 2019), with incubation on ice for 1 h followed by centrifugation at 4 °C for 20 min at 20,000 *g* using an Eppendorf Centrifuge 5430R microcentrifuge (Eppendorf Canada Ltd, Mississauga, ON, Canada), supernatant combined and either used same day or stored at –80 °C.

### 2.3 B vitamin standards

Non-labelled folate (B9) standards, 5,6,7,8-tetrahydrofolate (THF), 10-formyl folic acid (10-FFA), 5-formyltetrahydrofolate (5-FTHF), 5-methyltetrahydrofolate (5-MTHF), and 5,10-methenyltetrahydrofolate (5,10-methenyl THF) were purchased from Shircks Laboratories (Jona, Switzerland). MeFox (pyrazino-*s*-triazine derivative of 4a-hydroxy-5-methyltetrahydrofolate) was purchased from Merck & Cie (Schaffhausen, Switzerland). Folic acid (B9), thiamine (B1), riboflavin (B2), nicotinic acid (B3), pantothenic acid (B5), biotin B7), pyridoxine (B6) and pyridoxal (B6) were purchased from Sigma-Aldrich (St. Louis, MO, USA). Nicotinamide (B3) and pyridoxamine (B6) were purchased from Fisher Scientific (Waltham, MA, USA).

Isotope-labelled internal standards, ^13^C_5_-folic acid, ^13^C_5_-5-MTHF and ^13^C_5_-5-FTHF were purchased from Merck Eprova (Schaffhausen, Switzerland). ^13^C_4_^15^N_2_-riboflavin, ^13^C_3_^15^N-nicotinamide, ^13^C_3_^15^N-pantothenic acid, D_4_-biotin, D_3_-pyridoxine and ^13^C_3_-thiamine were purchased from Cambridge Isotope Laboratories (Tewksbury, MA, USA).

Stock solutions (1 mg/mL) and working solutions (0.1 mg/mL or 0.5 mg/mL) of B9 vitamers standards were prepared as previously described (Zhang *et al*., 2019), in 50 mM potassium phosphate buffer with 1% sodium ascorbate, 0.2% DTT and 0.2% MCE. 5,10-methenyl THF was prepared in pH 5 phosphate buffer, and all other B9 standards were prepared in pH 7 buffer. For the other B vitamins, stock solutions (1 mg/mL) and working solutions (0.1 mg/mL or 0.5 mg/mL) were prepared in water (Nurit *et al*., 2015). All solutions were aliquoted and stored at –80 °C prior to use.

### 2.4 B vitamin extraction

The extraction protocol used was similar to that described in Zhang *et al*., 2021. In brief, 30 mg (± 1.0 mg) finely ground seed powder was weighted into 2 mL microvials (Sarstedt, Germany). Method blank vials containing no seed powder were also prepared. To each sample vial, 170 μL of extraction buffer (pH 6.0, 50 mM MES containing 1% w/v sodium ascorbate and 0.2% v/v MCE) and 450 μL of extraction buffer containing nine stable isotope labelled internal standards (Section 2.3) was added and vortexed. Samples were heated for 5 min at 100 °C with shaking at 1200 rpm using a Vortemp 56 shaking incubator (Montreal-Biotech Inc., Kirkland, QC, Canada). Samples were cooled on ice. To the cooled mixture, 80 μL each of AP and BGL and 120 μL charcoal-treated rat serum were added, resulting in a final extraction volume of 900 μL. Samples were vortexed and incubated at 37 °C and 1200 rpm for 4 h in the Vortemp 56 shaking incubator. Samples were cooled on ice and centrifuged at 4 °C and 16,600 *g* for 30 min using an Eppendorf 5430R microcentrifuge. From the supernatant, 200 μL were transferred to 10 kDa MWCO filters (PALL, New York, NY, USA) and centrifuged as above. Filtrate was transferred to glass inserts inside amber vials for LC-SRM analysis (Agilent Technologies, Santa Clara, CA, USA). We analyzed six biological replicates for each 2018 genotype, tissue type, and growth site, and three biological replicates for each 2020 genotype and tissue type. Extractions from each biological replicate were injected twice for LC-MS.

### 2.5 Ultra-performance liquid chromatography-selected reaction monitoring mass spectrometry (UPLC-SRM MS) analysis

B vitamin separation and detection was performed using a Thermo Fisher Vanquish UPLC coupled with a TSQ Altis triple quadrupole mass spectrometer operated in positive electrospray ionization (ESI) mode (San Jose, CA, USA). Fragmentation of individual B vitamin standards was optimized by direct infusion, in a solvent of 50% acetonitrile containing 0.1% acetic acid. Details regarding SRM transitions and other parameters are discussed in Zhang *et al*., 2021.

An Agilent ZORBAX SB-Aq column (100 mm x 2.1 mm, 1.8 μm particle size) was used for B vitamer separation, with a SB-Aq guard column (5 mm x 2.1 mm, 1.8 μm particle size) used for sample loading and enrichment. The mobile phase consisted of 10 mM ammonium acetate, 0.1% acetic acid in either water (solvent A) or MeOH (solvent B). The flow rate was set to 0.3 mL/min and the gradient initially set to 95% solvent A and maintained for 30 s, the elution during this period was directed to waste through a divert valve. Solvent A was decreased to 85% from 30 s to 36 s and ramped to 10% until 3 min and maintained until 4 min. The composition was then changed back to 95% solvent A from 4 min to 4.1 min and maintained until 5 min to equilibrate the column. Elution from the last 0.9 min was directed to waste. The column compartment was maintained at 25 °C and the autosampler at 6 °C. The light in the autosampler was turned off, and the autosampler door was shielded. The mass spectrometer parameters were electrospray voltage 3000 V, vaporizer temperature 350 °C, sheath gas pressure 60, auxiliary gas pressure 10, sweep gas 1, and capillary temperature 325 °C. Six biological replicates were injected per 2018 genotype, three biological replicates were injected per 2020 genotype, and each biological replicate was injected twice.

### 2.6 Calibration curve and concentration determination

Nine labelled internal standards, ^13^C_3_-thiamine, ^13^C_4_^15^N_2_-riboflavin, ^13^C_3_^15^N-nicotinamide,^13^C_3_^15^N-pantothenic acid, D_3_ -pyridoxine, D_4_ -biotin, ^13^C_5_ -folic acid, ^13^C_5_ -5-FTHF, and ^13^C_5_ -5-MTHF were used for quantification, and calibration curves were prepared using sixteen B vitamin standards; thiamine, riboflavin, nicotinic acid, nicotinamide, pantothenic acid, pyridoxal, pyridoxine, pyridoxamine, biotin, folic acid, THF, 5-FTHF, 5-MTHF, 5,10-methenyl THF, 10-FFA, and MeFox. ^13^C_3_-thiamine was used to normalize thiamine; ^13^C_4_ ^15^N_2_ -riboflavin was used to normalize riboflavin; ^13^C_3_^15^N-nicotinamide was used to normalize nicotinic acid and nicotinamide; ^13^C_3_^15^N-pantothenic acid was used to normalize pantothenic acid; D_3_-pyridoxine was used to normalize pyridoxal, pyridoxine, and pyridoxamine; D_4_-biotin was used to normalize biotin; ^13^C_5_-folic acid was used to normalize folic acid, THF, and 10-FFA; ^13^C_5_-5-FTHF was used to normalize 5-FTHF; ^13^C_5_-5-MTHF was used to normalize 5-MTHF, 5,10-methenyl THF, and MeFox.

The concentration range of each standard was determined in initial experiments. An eight-point calibration curve was used for B vitamin quantification. A method blank including the internal standards was used for background signal quantification. A 1/x weighed linear regression was used for all vitamers as this provided good linearity and accurate measurement (Jha *et al*., 2015; Zhang *et al*., 2018a, 2019).

Due to vitamer instability, calibration curve solutions were prepared same day as the samples and were injected at the beginning, middle, and end of the queue. The peak area ratios of the target vitamer compared with the corresponding internal standard were used to build the calibration curve.

### 2.7 Statistical Analysis and Data Visualization

An ANOVA was performed using a mixed model analysis with the “lmer” function of the “lme4” package in R (R Development Core Team, 2019). Data met the assumptions of an ANOVA. The combination of year and location were considered an environment (site-year). The genotype and tissue type were treated as fixed effects, while site-year and replications (nested within site-years) were considered as random effects. To determine significant differences among tissue types and genotypes, means were separated using the “TukeyHSD” (Tukey Honest Significant Difference) function of the “mulcompView” package in R (Graves *et al*., 2019). Data was visualized using the ggbiplot function in R (Wickham, 2016).

## 3. Results

### 3.1 B vitamin differences by growth year for 2018 and 2020 samples

There were significant differences in B vitamin concentrations among lentil lines grown in both 2018 and 2020 (CDC Greenstar, IG72643, Eston, IG72623, Lupa, and BGE 016880) for both tissues examined (**Figure 1)**. Total B vitamin concentrations in cotyledon samples were lower in samples from 2018 versus samples from 2020, while total B vitamin concentrations in seed coats showed greater variability and a larger range of concentrations in samples from 2018, while samples from 2020 displayed more consistent total concentrations.

**Figure 1.**
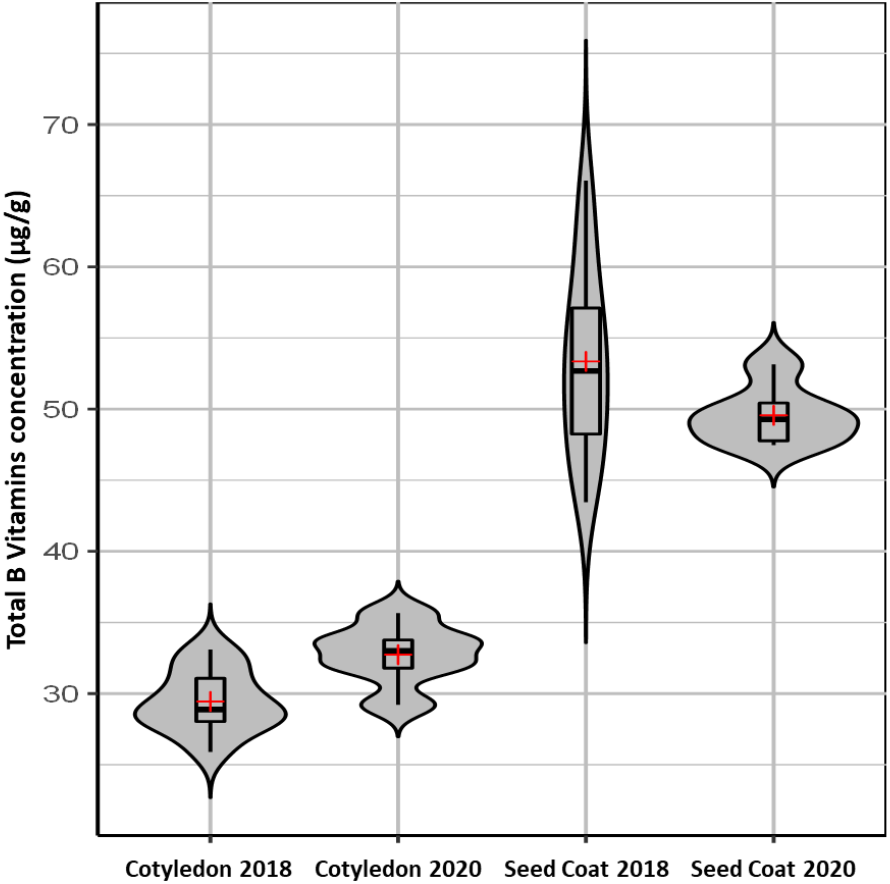
Total B Vitamins concentration (μg/g) in cotyledon and seed coat of lentil seeds of 37 *Lens* accessions grown during 2018 and 2020 in Saskatchewan Canada. TukeyHSD: 6.57.

The 2018 samples had lower measured concentrations of B1 and B3 in both tissues, B2 and B9 in seed coats, and B5 in cotyledons than in 2020. The 2018 samples had higher measured concentrations of B5 and B6 in seed coats (**Supplemental Table S1, Supplemental Table S2)**. No significant differences in concentrations were determined between 2018 and 2020 samples for B2, B6, and B9 in cotyledons. Concentrations of B7 were below the limit of quantification across the samples examined, and so their values have been excluded from these analyses. Overall, cultivars and wild accessions exhibited similar total B vitamins concentrations between 2018 and 2020 samples (**Supplemental Table S1**).

### 3.2 Tissue specific B vitamin concentrations in lentil

Significant differences in B vitamin concentrations were observed across tissue types in the set of thirty-four cultivars and three wild accessions grown in 2020 (**Figure 2**; **Supplemental Table S2; Supplemental Table S3**). For equal tissue weights, cotyledons had higher concentrations of vitamins B1 and B3, while seed coats had higher concentrations of B2, B5, B6 and B9. Cotyledon color (orange, yellow or green) did not have an effect on the concentration of any of the B vitamins. The distribution of B vitamins in both tissue types, followed the same trend in the wild accessions compared to cultivars, with the exception of the wild accession IG 72623 that had higher concentration of vitamin B1 in the seed coat compared to the cotyledon (**Supplemental Table S3**).

**Figure 2.**
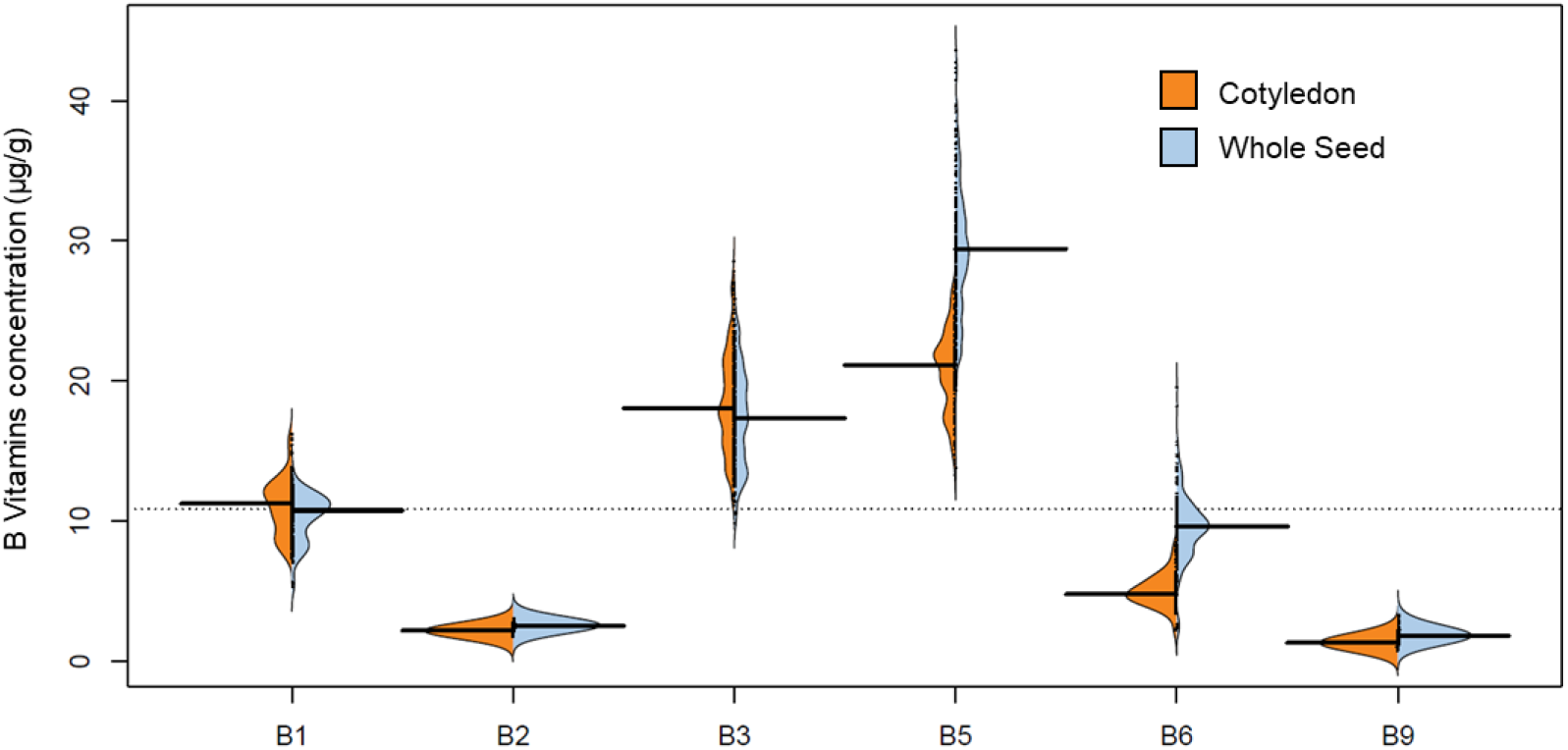
Values of B Vitamins concentration (μg/g) in cotyledon and whole seed of lentil seeds of 37 *Lens* accessions grown in 2020 in Saskatchewan Canada. TukeyHSD B1: 0.05, B2: 0.02, B3: 0.21, B5: 0.55, B6: 0.24, B9:0.02.

### 3.3 Genotype specific B vitamin concentrations in lentil

There were significant differences in vitamin concentrations among genotypes (**Figure 3**; **Supplemental Table S3**). Concentrations are reported as μg B vitamin per g lentil tissue flour. Vitamin B1 had concentrations ranging from 7.23 (ILL 9) to 12.98 (VIR-421) in whole seeds, 1.36 (IG 72623) to 15.99 (CDC Redberry) in cotyledons, and 1.73 (BGE 016880) to 8.80 (Guarena) in seed coats. Vitamin B2 had concentrations ranging from 2.16 (ILL 7727) to 2.95 (ILL 618) in whole seeds, 1.76 (ILL 1704-1) to 2.72 (IG 72623) in cotyledons, and 2.67 (ILL 1704-1) to 4.50 (BGE 016880) in seed coats. Vitamin B3 had concentrations ranging from 12.48 (Shasta) to 23.39 (CDC Milestone) in whole seeds, 11.21 (IG 72643) to 26.81 (CDC Redberry) in cotyledons, and 4.73 (ILL 1704-1) to 16.98 (CDC Robin) in seed coats. Vitamin B5 had concentrations ranging from 21.62 (ILL 4609) to 43.96 (CDC Redberry) in whole seeds, 7.05 (IG 72623) to 25.47 (4656-3) in cotyledons, and 18.55 (CDC Greenstar) to 60.85 (CDC Redberry) in seed coats. Vitamin B6 had concentrations ranging from 2.51 (ILL 9) to 16.35 (CDC Redberry) in whole seeds, 1.72 (IG 72623) to 7.95 (Guarena) in cotyledons, and 2.01 (ILL 9) to 39.42 (CDC Redberry) in seed coats. Vitamin B9 had concentrations ranging from 1.12 (Lupa) to 2.77 (ILL 618) in whole seeds, 0.89 (2275-15) to 2.87 (IG 72643) in cotyledons, and 1.30 (Shasta) to 8.86 (BGE 016880) in seed coats.

**Figure 3.**
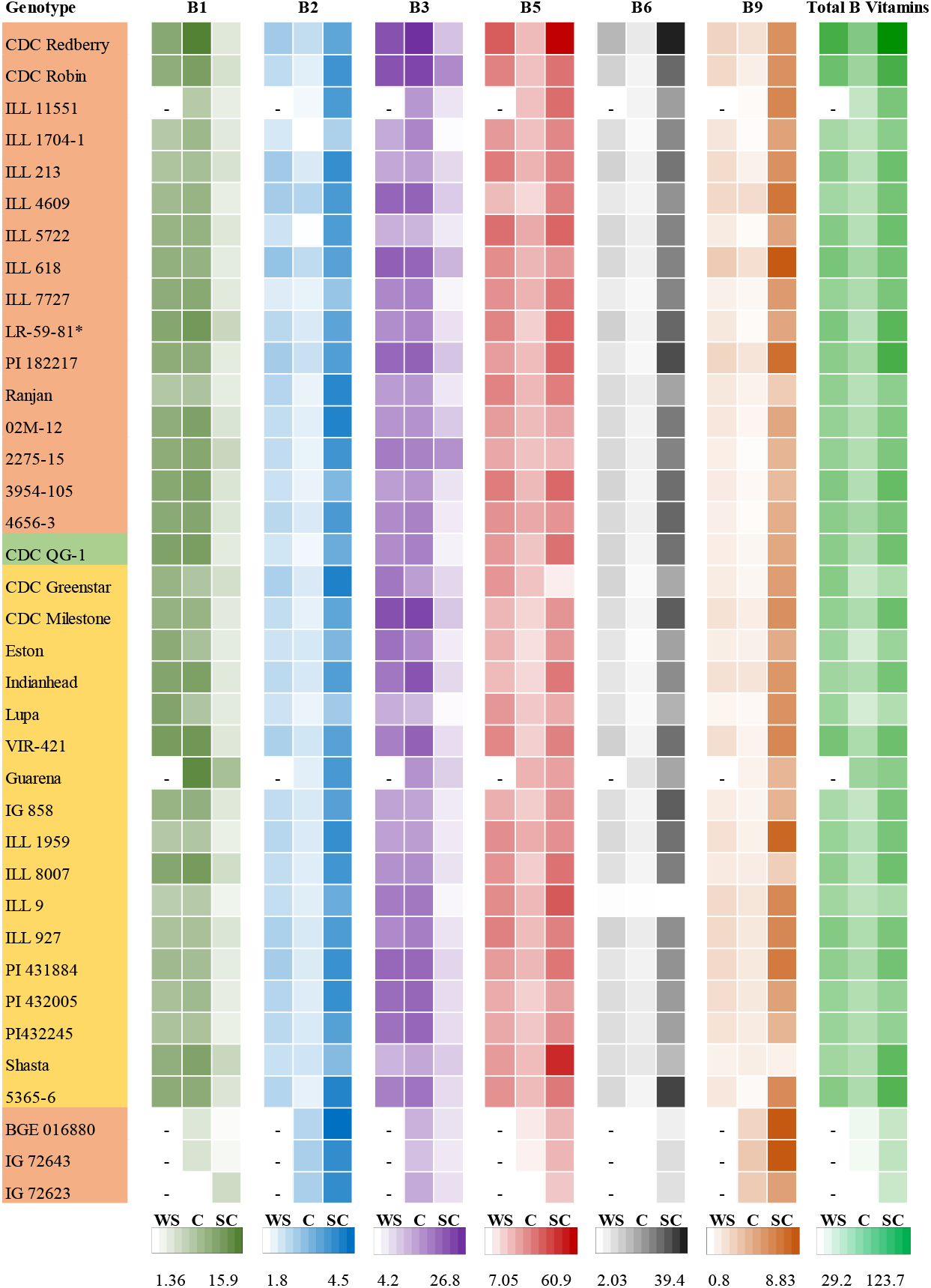
Values of B Vitamins concentration (μg/g) in whole seed, cotyledon, and seed coat of lentil seeds of 37 *Lens* accessions grown in 2020 in Saskatchewan Canada. TukeyHSD B1: 0.54, B2: 0.24, B3: 3.28, B5: 5.93, B6: 2.54 and B9: 0.26. Colors on genotypes indicate market classes: red, yellow and green. “–” indicates absence of data.

CDC Redberry consistently showed high vitamin concentrations across the B vitamin families, possessing the: highest cotyledon B1 concentration, second highest whole seed B2 concentration, second highest whole seed B3 concentration, highest whole seed B5 concentration, highest whole seed B6 concentration, and second highest whole seed B9 concentration.

Examining the concentrations of B vitamin families across genotypes also allowed for the identification of genotypes with tissue specific vitamin distribution unlike the others. One of the best examples is the cultivar Shasta. For vitamin B2, Shasta had the eighth lowest concentration for whole seed, but the concentration in cotyledon was eighth highest, and the concentration in seed coat was fourth lowest. For vitamin B5, Shasta whole seed concentration was twelfth lowest, while the concentration in seed coat was second highest. For vitamin B6, Shasta whole seed concentrations were twelfth lowest, while the concentration in cotyledon was second highest, and the concentration in seed coat was fifth lowest. For vitamin B9, Shasta whole seed concentration was second lowest, while the concentration in cotyledon was sixteenth highest, and the concentration in seed coat was the lowest.

The three wild accessions, IG 72643, IG 72623, and BGE 016880, typically grouped together based on vitamin concentrations in the cotyledons. They possessed the lowest three B1, B5 and B6 concentrations, the three highest B9 concentrations, and among the highest B2 concentrations (**Figure 3**).

## 4. Discussion

Establishing the complete B vitamins profile of a large set of lentils was necessary to determine the potential of breeding to increase B vitamins concentration. It was also important to explore any potential differences between cultivars and wild accessions, as part of the effort to understand all the potential contributions, positive or negative, from wild related species to the lentil crop. By analyzing whole seeds and cotyledons only, we gained a better understanding of the impact of removing seed coat in the concentration of B vitamins. The observed concentrations of B vitamins across analyzed lentil genotypes, reveals a range of important observations for production and consumption.

Initial comparison between samples harvested in 2018 and 2020 revealed the degradation or interconversion of B vitamins during storage. As the majority of significant B vitamin changes between sample years occurred in the seed coat, we suspect these concentrations are the result of degradation, interconversion, or consumption of these B vitamins during storage conditions. This confirms that MeFox is predominantly present in the seed coat, as previously proposed by Zhang *et al*. (Zhang *et al*., 2019).

We also detected significant increases in MeFox concentrations in the seed coats of 2018 samples (**Supplemental Table S1**). It is likely that oxidation during the storage process was one of the major contributors to the observed changes. As MeFox is a metabolically unavailable oxidation product of members of the B9 family, this observation supports the possibility of storage-based oxidation of B vitamins as a cause of some of the observed changes in vitamin concentrations between 2018 and 2020 samples.

This result demonstrates that longer periods of seed storage prior to analysis can result in changes to the overall B vitamin concentrations detected. As lentils may be stored for significant periods of time prior to consumption, be it in storage bins following harvesting, shipping containers for transport, or packaging on store shelves before being purchased by customers, the actual amount of vitamins remaining at time of purchase available for consumers could be significantly different from those calculated during earlier analyses. The lentils we examined were stored as whole seeds, so the seed coat may have shielded the cotyledon from oxidation. However, with many lentils commercially sold as splits, dehulled cotyledons may be stored for a long time which would expose them to potential oxidation and decreased vitamin availability. One possibility could be for different genotypes to have different roles in consumption, with those showing higher whole seed concentrations being most beneficial for eating whole, while those with higher concentrations in the cotyledon would prove superior for preparing split lentils or dehulled lentil flour.

For larger analysis alongside the initial cultivars, only the samples from 2020 were selected, in order to minimize impact of different storage periods on detected vitamin concentrations.

Our analysis included dramatically more genotypes of lentils than previous analyses, especially for vitamins B1, B2, B3, B5, and B6, which only had minimal coverage previously (**Table 2**). This analysis also provides a much more extensive look at the range of vitamin concentrations observed across genotypes. For whole seeds between the lowest and highest concentrations we observed; B1 had 1.80 fold increase, B2 had a 1.37 fold increase, B3 had a 1.87 fold increase, B5 had a 2.03 fold increase, B6 had a 6.51 fold increase, and B9 had a 2.47 fold increase. This demonstrates the significant role genotype plays in lentil B vitamin concentrations and helped us identified those worthy of further study and breeding for increased B vitamin content.

**Table 2:**
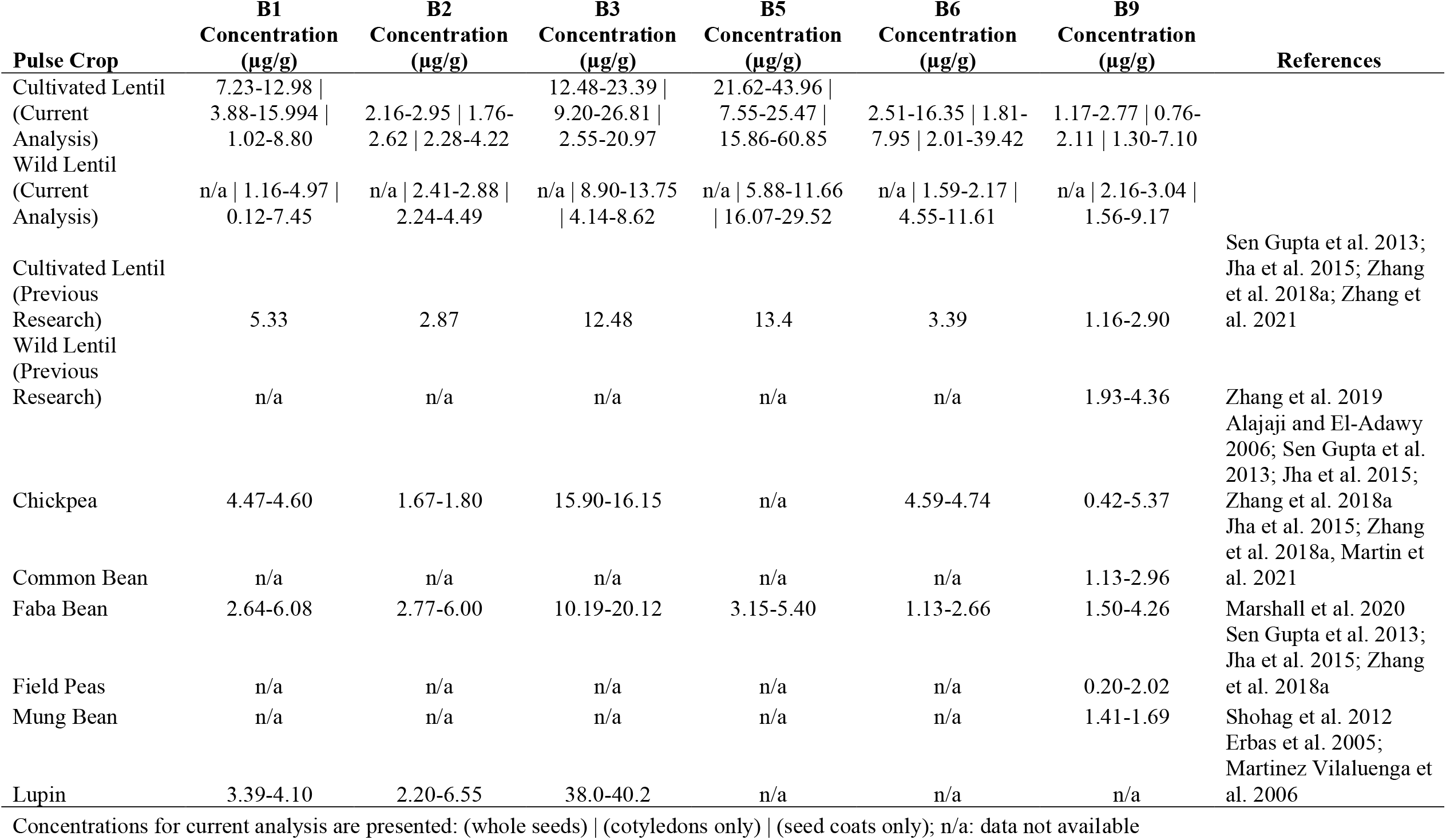
B vitamin concentrations across pulse species.

Given that CDC Redberry was among the cultivars with superior values for most B vitamins, studying populations that have CDC Redberry as a parental line may assist in understanding the inheritance on B vitamin concentration, and if its progeny will show higher concentrations of any of the B vitamins (transgressive segregation).

The observations in Shasta are also particularly interesting as Shasta is a zero-tannin line.

Tannins are compounds originally identified for their usefulness in tanning hides into leather through their crosslinking of proteins and have also been associated with a variety of positive and negative health effects in human consumption (Chung *et al*., 1998). As Shasta had lower seed coat concentrations for several B vitamins, it is possible that tannins may help protect B vitamins from degradation when exposed to either oxygen or bright light. However, as the Shasta lentils had higher cotyledon concentrations for B2, B6, and B9, it is possible that not having to produce tannins results in additional resources available for synthesis of B vitamins.

With the greater depth of B vitamin analysis across many lentil genotypes, it also allows for further comparisons between the concentrations present in lentils and other pulses (**Table 2**). Chickpea shows concentrations of lower B1 and B2, B3 concentrations near the middle range of lentil concentrations, B6 concentrations near the lower range of lentil concentrations, and B9 concentrations lower than lentil at their lower concentrations and higher than lentil at their higher concentrations, with no B5 concentrations determined (Alajaji and El-Adawy, 2006; Sen Gupta *et al*., 2013; Jha *et al*., 2015; Zhang *et al*., 2018a). Common beans have B9 concentrations near or slightly above the upper concentrations of cultivated lentil lines (Sen Gupta *et al*., 2013; Jha *et al*., 2015; Zhang *et al*., 2018a; Martin *et al*., 2021). Our previous analysis of faba bean had B1 concentrations lower than lentil, B2 concentrations at or above the highest concentrations of lentil, B3 concentrations similar to lentil, B5 concentrations lower than lentil, B6 concentrations below or at the lower concentrations of lentils, and B9 concentrations that range from within the range of lentil concentrations to higher concentrations than lentil (Marshall *et al*., 2020). Field pea has B9 concentrations lower than, or in the middle of lentil (Sen Gupta *et al*., 2013; Jha *et al*., 2015; Zhang *et al*., 2018a). Mung beans have B9 concentrations in the lower range of lentil (1.41-1.69 μg/g). Lupin have B1 concentrations lower than lentil, B2 concentrations ranging across and higher than lentil, and B3 higher than lentil (Erbas *et al*., 2005; Martinez Vilaluenga *et al*., 2006).

The other previous analysis of wild lentils (Zhang *et al*., 2019) only examined B9, and also found higher concentrations of vitamin B9 in wild accessions compared to cultivated lentils. These results demonstrate that in addition to the disease, pest, and abiotic stresses, wild accessions also have the potential to contribute with higher B2 or B9 concentrations to cultivated lentils. However, the potential to yield higher concentrations in those vitamins would need to be weighed against the potential decrease in other vitamin concentrations. The wild accessions evaluated are from the species *L. orientalis* (BGE 016880 and IG 72643) and *L. odemensis* (IG 72623). The complete B vitamins profile needs to be studied on individuals from the other four wild species related to the lentil crop to determine if there are other sources useful for increasing these in cultivated lentil.

## 5. Conclusion

This research is, to our knowledge, the first large scale quantification and comparison of B vitamin levels in a globally diverse set of lentil genotypes, including cultivated and wild accessions, and analyzing different seed tissues. It demonstrates the impact of storage on B vitamin concentrations, with the increased MeFox levels suggesting oxidation is a major contributing factor, which in turn has implications for reporting B vitamin levels on lentil products stored for extended periods of time prior to consumption. Tissue specific distribution of B vitamin families was observed, with cotyledons containing higher concentrations of B1 and B3, and seed coats containing higher concentrations of B2, B5, B6, and B9. Most red lentils sold commercially have their seed coats removed, resulting in all of the vitamins present in the seed coat being lost. Consumption of whole lentils can avoid these losses and offer consumers lentil products higher in these B vitamins. In addition, we have demonstrated the wide range of B vitamin concentrations present within the diverse lines of lentils analyzed. With significant concentration differences across lines, B vitamin concentrations are worth monitoring while breeding for lines bearing advantageous traits to see if higher vitamin yields can also be obtained. As the wild lines examined showed higher B2 and B9 concentrations than cultivated lines, vitamin concentrations could be another trait that could be introgressed from wild related species to cultivated lines along with the usual stress and disease resistance traits.

## List of Abbreviations

MCE: 2-Mercaptoethanol
MES: 2-(N-morpholino)ethanesulfonic acid
AP: acid phosphatase
BGL: β-glucosidase from almonds
DTT: dithiothreitol
THF: 5,6,7,8-tetrahydrofolate
10-FFA: 10-formyl folic acid
5-FTHF: 5-formyltetrahydrofolate
5-MTHF: 5-methyltetrahydrofolate
5,10-methenyl THF: 5,10-methenyltetrahydrofolate
MeFox: pyrazino-*s*-triazine derivative of 4a-hydroxy-5-methyltetrahydrofolate
ESI: electrospray ionization

## Declaration of Competing Interest

The authors declare that they have no known competing financial interests or personal relationships that could have appeared to influence the work reported in this paper.

## Acknowledgements

This work was supported by the ‘Enhancing the Value of Lentil Variation for Ecosystem Survival (EVOLVES)’ project funded by Genome Canada [grant: LSP18-16302] and managed by Genome Prairie. We are grateful for the matching financial support from Western Grains Research Foundation [grant: GC1903], the Government of Saskatchewan [grant: 20200026, and the University of Saskatchewan. We thank the Pulse Crop Field Lab technical staff at the University of Saskatchewan for their assistance with plant and seed production and management. We thank Dr. Albert Vandenberg at the University of Saskatchewan for the use of his TSQ Altis mass spectrometer for sample analysis. We thank Dr. Haixia Zhang for methodology optimization and mass spectrometry technical expertise and assistance.

## Appendix A. Supplementary data

Supplementary data associated with this article are available.

## Data availability Statement

- The data supporting this study are available online (https://knowpulse.usask.ca/experiment/phenomics/B-vitamin-quantification-in-wild-and-cultivated-lentil-seed-tissues-using-ultra-performance-liquid-chromatography-selected-reaction-monitoring-mass-spectrometry)

## Notes

### Competing Interest Statement

The authors have declared no competing interest.

